# Demonstration of an Integrated Process for Manure-Based Nitrogen Recovery Using the Biopolymer Cyanophycin

**DOI:** 10.64898/2026.07.29.741614

**Authors:** Kevin Fitzgerald, Hang Dong, Edward Apraku, Md Aminul Islam Prodhan, Dylan Hakken, George Wells, William Tarpeh, Keith Tyo

**Author notes:** Corresponding author: Keith Tyo. Currently at: School of Civil and Environmental Engineering, Georgia Tech Shenzen Institute, Tianjin University, Shenzhen, Guangdong 518055, China (Hang Dong).

## Abstract

The trend towards concentrated animal feeding operations (CAFOs) has served to concentrate not only livestock animals but the waste they produce to comparatively smaller areas. The point-source nature of this waste is an opportunity for the recovery and valorization of the nitrogen therein. Such a process would be viable on small to intermediate scales and require minimal inputs at the CAFO. In this study, we demonstrate the potential of the biopolymer cyanophycin to serve as a medium for manure-nitrogen recovery. In the first step, genetically modified strains of *Escherichia coli* produce intracellular cyanophycin from mock manure hydrolysates. Next, cyanophycin is recovered from microbial biomass via acid solubilization and base precipitation using electrochemically generated acids and bases.

Finally, to improve both the yield and recoverable fraction of cyanophycin produced, we leverage the tunability of our genetically engineered system to probe the impacts of cyanophycin synthetase solubility, N-domain activity, and cyanophycin molecular weight on cyanophycin recoverability. Collectively, this work serves as a proof of concept for nitrogen recovery from agricultural waste, aligning with global sustainability initiatives.

## 1. Introduction

Organic carbon-, nitrogen-, and phosphorus-rich wet waste streams are produced in substantial quantities across both urban and agricultural sectors and are increasingly recognized as valuable feedstocks for chemical and biological valorization (Milbrandt et al., 2018),(Philbrick et al., 2017). Livestock waste constitutes a substantial source of nitrogen in the United States, annually accounting for 8.1 million tons of nitrogen from 1.4 billion tons of manure (Pagliari et al., 2020). Concurrently, contemporary livestock practices have trended towards increased animal stocking densities, exemplified by the proliferation of concentrated animal feeding operations (CAFOs) (Copeland, 2010). As a byproduct of consolidating livestock to smaller areas, CAFOs inherently localize waste generation. Unlike non-point source emissions, this point source waste is increasingly viewed as a promising source of substrate, as it can be more readily collected, treated, and converted into value-added products.

Manure is currently treated as a waste stream and represents significant environmental and logistical challenges (Copeland, 2010). CAFO-derived manure is typically collected and concentrated in open-air lagoons or underground pits, diluted with water to facilitate transport, and then, ideally, land-applied as fertilizer. However, manure is often produced in volumes that exceed the nutrient assimilation capacity of nearby farmland. Once this capacity is reached, additional manure must be transported greater distances prior to application, increasing costs. An upper bound for nitrogen concentration in manure is approximately 0.5% by weight, up to two orders of magnitude lower than that of common synthetic fertilizers like urea (46.7% N).

Given manure’s low nitrogen concentration on a wet-weight basis, these transportation costs can significantly undermine its commercial viability as a nitrogen fertilizer (Ribaudo et al., 2003). As a result, full-compliance with EPA mandates is estimated to incur additional manure-transport related costs on the order of two billion dollars annually (Ribaudo et al., 2003). For this reason, the conversion of manure to an alternative, concentrated form more suitable for nitrogen recovery, storage, and utilization offers distinct advantages over prevailing practices.

The biological upscaling of waste via nutrient concentration is not without precedent; enhanced biological phosphate removal (EBPR) processes, for instance, rely on the concentration of environmental phosphorus into intracellular polyphosphate prior to downstream recovery as struvite. Alternative nitrogen recovery methods are predominantly physicochemical in nature but suffer from pragmatic limitations: air stripping, for instance, requires high energy inputs at elevated recovery levels, while adsorption-based strategies are prone to fouling in agricultural environments (Pandey & Chen, 2021),(Zhou et al., 2023),(Song et al., 2021),(Al-Juboori et al., 2022). Despite the strong precedent of biological solutions for nitrogen *removal* (i.e., anammox, nitrification–denitrification), biological solutions for nitrogen *recovery* remain comparatively underexplored, particularly in the context of agricultural waste.

## 2. Background

Cyanophycin, or cyanophycin granule peptide (CGP), is an intracellular polymer with properties amenable to the microbial recovery of waste nitrogen (Berg et al., 2000).

Cyanophycin is a non-ribosomal peptide featuring a poly-L-aspartic acid backbone from which L-arginine or L-lysine attach via isopeptide bonds. As a function of this composition, cyanophycin has a relatively high nitrogen content of ∼23% (w/w). The polymerization reaction is enabled by the activity of a single enzyme, cyanophycin synthetase (CphA1). Reflecting its broad biological compatibility, cyanophycin synthetase expression and cyanophycin production has been demonstrated in several industrially relevant organisms, including *Escherichia coli*, *Corynebacterium glutamicum*, *Saccharomyces cerevisiae*, *Pseudomonas putida*, *Pichia pastoris*, *Ralstonia eutropha*, and *Rhizopus oryzae* (Du et al., 2019),(Xiao et al., 2017),(Wiefel et al., 2019). Previous work has also demonstrated cyanophycin’s efficacy as a nitrogen recovery vehicle for an ammonium-rich synthetic urine medium (Canizales et al., 2023).

In the context of animal agricultural, purified cyanophycin shows promise as a dietary supplement compatible with contemporary feeding practices. Past isolation of cyanophycin-degrading bacteria from the gut flora of cows, pigs, chickens, and fish suggests that the polymer could serve as a broadly digestible source of amino acids (Sallam & Steinbüchel, 2009). Cyanophycin is particularly promising in this context as a source of lysine, an essential amino acid for swine and poultry that is often deficient in typical feeds without explicit supplementation (Hasan et al., 2020),(Macelline et al., 2021).

In addition to its high nitrogen content, cyanophycin is compatible with relatively simple extraction techniques due to its pH-responsive solubility profile. Both arginine and lysine side chains contribute to the zwitterionic nature of cyanophycin. At neutral pH their respective guanidinium and ε-amino groups are protonated while their alpha carboxyl groups are deprotonated. These opposing charges enable the intermolecular attraction of disparate cyanophycin strands to create “insoluble” cyanophycin granules in pH-neutral conditions.

Under extreme pH conditions, however, one set of charges disappears: at high pH the guanidinium groups (pKa = 12.5) are mainly deprotonated and uncharged whereas at low pH the *a*-carboxyl group (pKa = 2.2) are mainly protonated and uncharged. This loss of one charge makes it no longer a zwitterion, reverting the aggregation. Many extraction protocols leverage this unique property (insoluble at neutral pH, but soluble at high/low pH) to separate cyanophycin from non-cyanophycin biomass (Steinle & Steinbüchel, 2010),(Frey et al., 2002),(Swain et al., 2023).

The efficacy of these simple extraction procedures for large-scale processing is undermined by the costs associated with the production, storage, and transport of the strong acids and bases required. Product purification can represent 20–40% of total production cost for large-scale fermentations, making the development of low-cost extraction strategies a critical factor for improving process economics (Straathof, 2011). An emerging alternative involves electrochemical generation of acids and bases through water splitting via oxidation and reduction reactions at electrodes or through water dissociation facilitated by bipolar membranes. This approach enables on-site, on-demand acid and base production to enable nitrogen recovery at CAFOs. Furthermore, the use of manure as a cost-negative substrate helps alleviate feedstock costs, which typically account for 15–60% of overall bioprocess expenditures (Straathof, 2011). These exploitations in substrate sourcing and product recovery can enhance the economic viability of cyanophycin-based nitrogen reclamation.

The work presented here aims to derisk the key technological advances necessary for the full realization of this proposed bioprocess. We first heterologously expressed three CphA1 homologs in *E. coli* and evaluated their capacity for cyanophycin production. Next, we demonstrated the recovery of nitrogen as cyanophycin from mock manure hydrolysate media. A small electrochemical reactor was then constructed to evaluate whether electrochemically generated acid and base could be used to control cyanophycin solubility for selective recovery. Finally, to guide future engineering strategies aimed at increasing total cyanophycin yield and improving the fraction recoverable via pH manipulation, we engineered several CphA1 mutants to explore how enzyme solubility and the CphA1 N-domain affect cyanophycin synthesis.

Together, these approaches establish a proof of concept for transforming waste nitrogen into a usable product at the site of manure generation.

## 3. Materials and Methods

### 3.1 Strains

Chemically competent *Escherichia coli BL21*(DE3) pLysE was purchased from Fisher Scientific (Waltham, MA) and used for all manure hydrolysate experiments. Chemically competent Arctic Express (DE3) *E. coli* was purchased from Agilent (Santa Clara, CA). Wild type *Acinetobacter baylyi* ADP1 was generously provided by Ellen Neidle (University of Georgia).

### 3.2 Plasmids and Cloning

The following cyanophycin synthetase (CphA1) homologs were used in this study: 6308CphA1 from *Synechocystis sp.* PCC6308, AbCphA1 from *Acinetobacter baylyi* ADP1, TmCphA1 from *Tatumella morbirosei*, and SuCphA1 from *Synechocystis* sp. UTEX2470. The protein sequence for 6308CphA1 was obtained from UniProt (accession number P56947), codon optimized for expression in *E. coli*, and synthesized via Integrated DNA Technologies (Coralville, IA). The gene for AbCphA1 was amplified directly from the genome of *A. baylyi* ADP1. The gene sequence for TmCphA1 and the N-domain of SuCphA1 as used previously (Sharon et al., 2021) was generously provided by Prof. Martin Shmeing (McGill University) and synthesized via Integrated DNA Technologies. Codon optimized nucleic acid sequences are available in SI Table 1.

**Table 1.** Plasmids used in this study.

| Plasmids | Description |
| --- | --- |
| pET28a-T7-mCherry | Non-cyanophycin accumulation control strain expressing red fluorescent protein |
| pET28a-T7-6308CphA1 | Cyanophycin synthetase CphA1 from *Synechosystis* sp. 6308 |
| pET28a-T7-AbCphA1 | Cyanophycin synthetase CphA1 from *Acinetobacter baylyi* ADP1 |
| pET28a-T7-TmCphA1 | Cyanophycin synthetase CphA1 from *Tatumella morbirosei* |
| pET28a-T7-MBP-6308CphA1 | Cyanophycin synthetase CphA1 from *Synechosystis* sp. 6308 tagged with maltose binding protein (MBP) fused to the N-terminus |
| pET28a-T7-SUMO-6308CphA1 | Cyanophycin synthetase CphA1 from *Synechosystis* sp. 6308 tagged with small ubiquitin-like modifier (SUMO) solubility tag fused to the N-terminus |
| pET28a-T7-Tm-6308CphA1 | Chimeric CphA1 fusion protein consisting of the N-terminal 149 amino acids from TmCphA1 fused to the remaining 712 amino acids of 6308CphA1. |
| pET28a-T7-Su-6308CphA1 | Chimeric CphA1 fusion protein consisting of the N-terminal 163 amino acids from SuCphA1 fused to the remaining 712 amino acids of 6308CphA1. |
| pET28a-T7-Ab-6308CphA1 | Chimeric CphA1 fusion protein consisting of the N-terminal 162 amino acids from AbCphA1 fused to the remaining 712 amino acids of 6308CphA1. |
| pET28a-T7-6308-AbCphA1 | Chimeric CphA1 fusion protein consisting of the N-terminal 162 amino acids from 6308CphA1 fused to the remaining 753 amino acids of AbCphA1. |

All gene inserts were integrated into pET28a(+) vectors via Gibson assembly using Gibson master mix (New England Biolabs, Ipswitch, MA) and transformed into chemically competent *E. coli* DH5*a*. Following selection with 25 μg/mL kanamycin monosulfate on LB-agar plates, transformants were screened via colony PCR and positive colonies were miniprepped and submitted for whole-plasmid sequencing. Plasmids with the correct sequence were transformed into chemically competent *E. coli* BL21(DE3) pLysE for all subsequent cyanophycin cultivation experiments.

N-domain replacements for 6308CphA1 were constructed using the first 149, 163, or 162 amino acids of TmCphA1, SuCphA1, or AbCphA1, respectively, fused to the last 712 amino acids of 6308CphA1. 6308-AbCphA1 used the first 162 amino acids of 6308CphA1 and the final 753 amino acids of AbCphA1. A description of all plasmids used in this study is shown below in Table 1. Sequences for all genetic parts can be found in SI Table 1.

### 3.3 Mock Manure Hydrolysate Medium

The primary components of the mock manure hydrolysate media were selected based on previous work using the reported hydrolysis products of dairy cattle manure (Chen et al., 2003). The hydrolysis products from the equivalent of 100 g dried manure was added to 1 L water to create a media with the following composition: 19.37 g/L D-glucose, 11.00 g/L D-xylose, 1.47 g/L L-arabinose, and 7.92 g/L casamino acids. Beyond manure hydrolysate components, the following media components were supplemented: 6.8 g/L sodium phosphate dibasic, 3.0 g/L potassium phosphate monobasic, 1.32 g/L ammonium sulfate, 476 mg/L sodium chloride, 240 mg/L magnesium sulfate, 17 mg/L calcium chloride, 6 mg/L iron (III) chloride, 6 mg/L manganese chloride, 6 mg/L copper (II) sulfate, and 6 mg/L zinc chloride. To probe the impact of nitrogen source on cyanophycin recovery, a high ammonium condition was made by replacing approximately 60 mM casamino acids (6.92 g/L) with 60 mM ammonium sulfate (7.93 g/L).

### 3.4 Culture Conditions for Cyanophycin Production

Individual colonies were selected from LB-agar plates and precultured overnight in 5 mL liquid LB medium at 30°C, 250 rpm. The following day, overnight cultures were added to either LB supplemented with 10 mM L-arginine (Fig. 1 and 2) or mock manure hydrolysate medium (Fig. 3) to a target OD600nm of 0.05 and grown at 30°C until the mid-exponential phase. At a target OD600nm of 0.5, cultures were transferred to 12°C and CphA1 expression was induced with 0.75 mM isopropyl β-D-1-thiogalactopyranoside (IPTG) after 10 min (SI Fig. 1).

**Figure 1.**
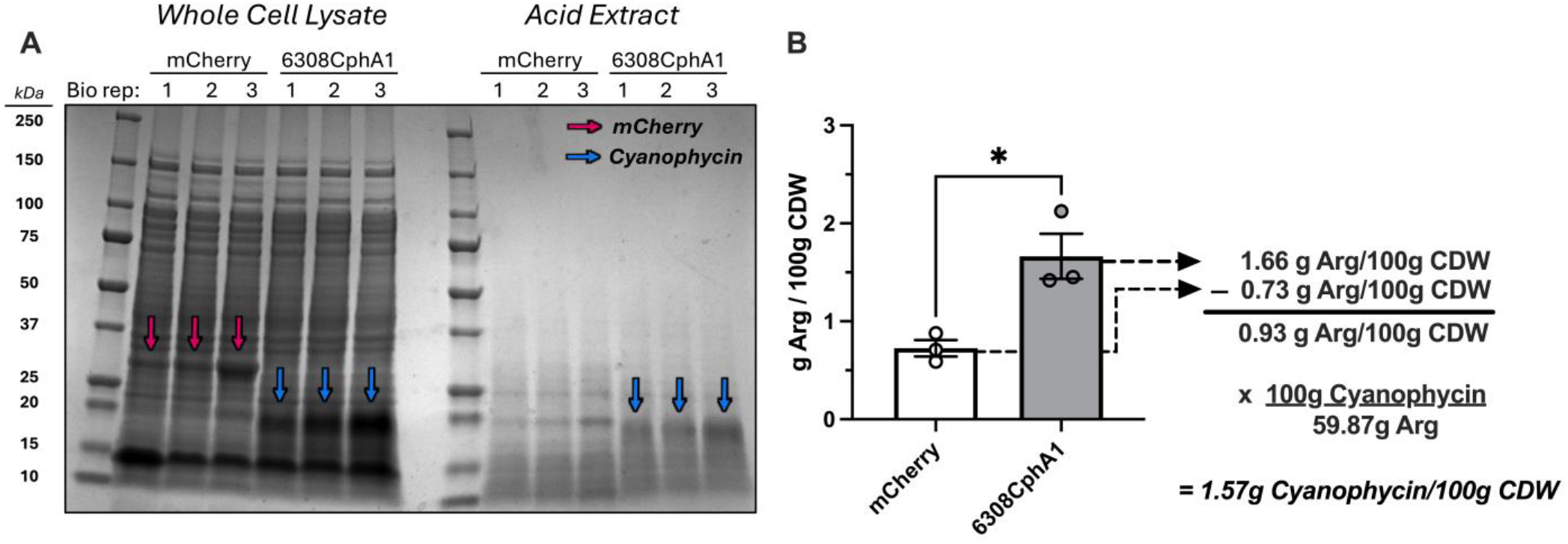
Cyanophycin visualization and quantification by correcting for background arginine. (A) SDS-PAGE gels visualizing cyanophycin in whole cell lysate and acid extract. mCherry (no cyanophycin control) and cyanophycin bands expected at 27 kDa and 8-25 kDa (Kwiatos & Steinbüchel, 2021), respectively. (B) Cyanophycin quantification workflow following acid extraction and arginine measurement via Sakaguchi reagent. The difference in specific arginine content between sample and mCherry negative control is attributed to cyanophycin. Cyanophycin content can then be estimated using an assumed mol% arginine. Arginine measurements were performed in technical duplicate and averaged for each biological replicate. *E. coli* culturing was performed using 50 mL LB medium supplemented with 10 mM arginine in 250 mL flasks. Error bars = SEM for biological triplicate. Asterisks indicate statistical significance compared to control (p < 0.05, Student’s t-test).

**Figure 2.**
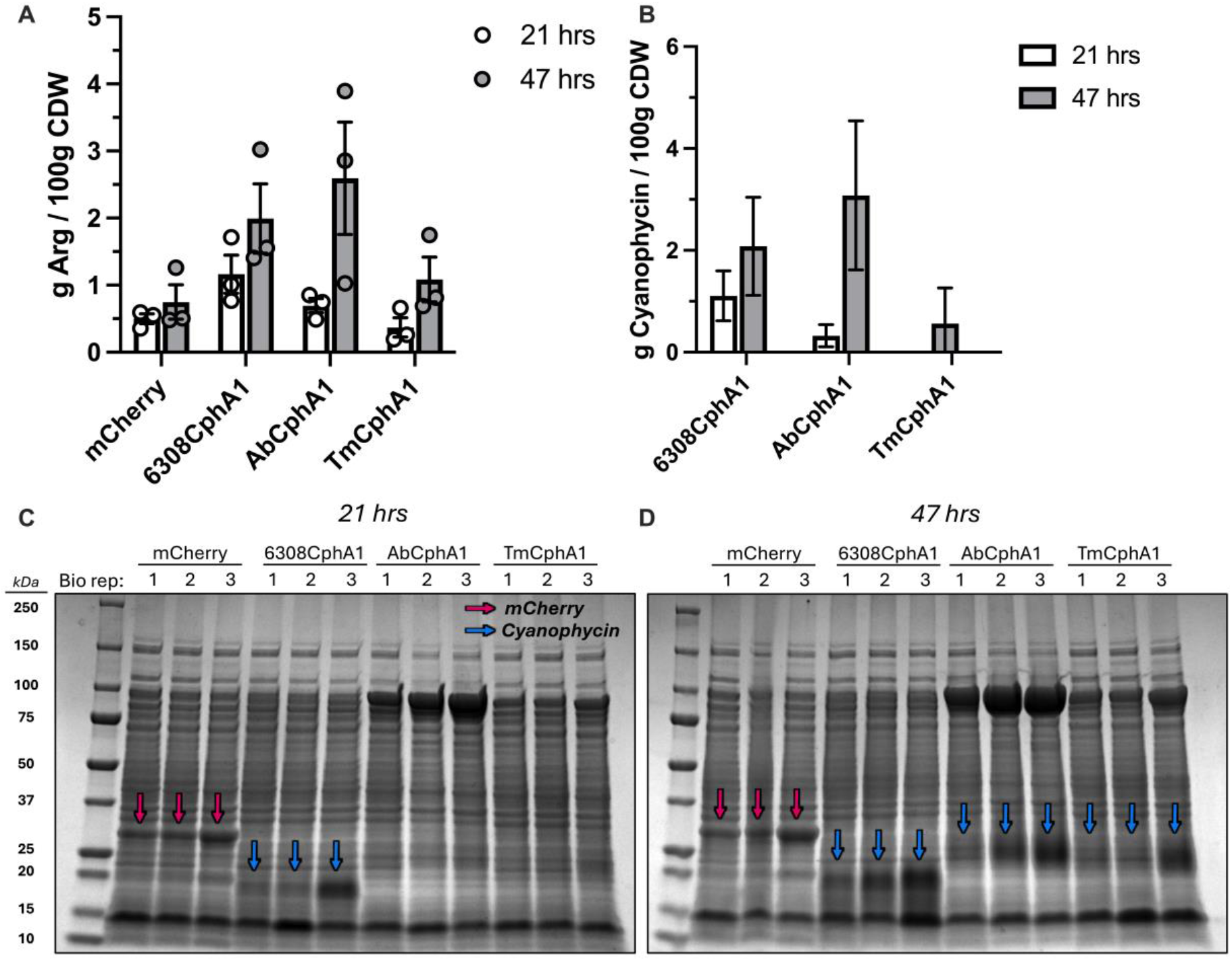
Cyanophycin synthetase homologs vary in apparent onset of polymer production, consistent with putative N-domain catalytic differences. (A) Biomass-normalized arginine content of acid extracts from *E. coli* strains expressing different CphA1 homologs or mCherry as a no-cyanophycin control. All strains were cultured in LB medium supplemented with 10 mM L-arginine. (B) Biomass-normalized cyanophycin titer in CphA1-expressing strains, estimated after subtraction of the no-cyanophycin control (mCherry) baseline. Data points yielding negative values (due to lower arginine content in the acid extract compared to no-cyanophycin control) were excluded (TmCphA1, 21 hrs) Whole cell lysates were visualized via SDS page at 21 hrs (C) and 47 hrs (D). Growth and fluorescence data shown in SI Fig. 4. Culturing was performed using 5 mL medium in 10 mL, 24-well plates with Breathe-Easy Sealing membranes (Millipore Sigma, Burlington, MA). Error bars = SEM for biological triplicate.

**Figure 3.**
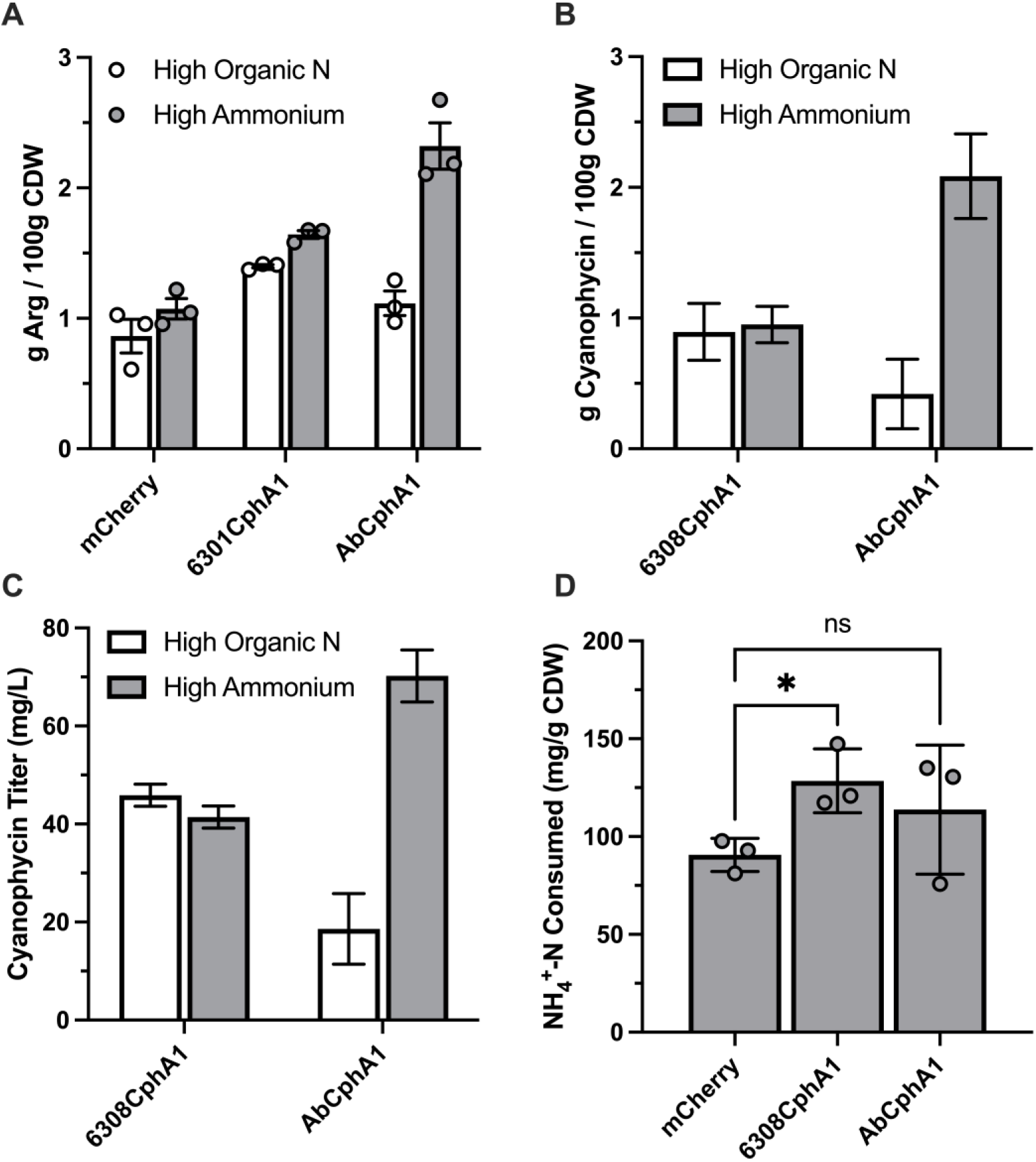
Nitrogen from mock manure hydrolysate media is efficiently recovered as cyanophycin. (A) Biomass-normalized arginine content of acid extracts from *E. coli* strains expressing different CphA1 homologs in mock manure hydrolysate media. (B) Biomass-normalized cyanophycin titers or (C) volumetric cyanophycin titers from CphA1-expressing strains, estimated after subtraction of the mCherry no-cyanophycin baseline. (D) Specific ammonium consumption in “High Ammonium” condition after 7 days of cultivation. Arginine measurements were performed in technical triplicate and averaged for each biological replicate. Culturing was performed using 50 mL medium in 250 mL flasks at 12 °C. Error bars = SEM for biological triplicate. Asterisks indicate statistical significance compared to control (p < 0.05, Student’s t-test).

### 3.5 Cyanophycin Purification

Cell pellets were collected via centrifugation at 5000 x g for 30 min at 4°C.

Pellets were first washed with 1 culture-volume ultrapure water (e.g. 50 mL water for 50 mL culture) to remove trace salts and arginine and then resuspended in 1/9th culture-volume ultrapure water. To lyse cells and solubilize intracellular cyanophycin, the washed cell suspension was adjusted to 0.1 M HCl by addition of 1 M HCl and incubated at room temperature with gentle inversion for 1 hr (SI Fig. 2). Following acid extraction, the remaining solids were separated from the supernatant via centrifugation and discarded. For downstream electrochemical solubility tests, the supernatant was pH-neutralized and the resulting precipitate was collected as insoluble cyanophycin (iCGP).

### 3.6 Electrochemical Acid and Base Generation

Acid and base production was achieved using a bipolar membrane which dissociates water into H⁺ and OH⁻ under an applied electric field. The bipolar membrane was integrated into a custom-built three chamber electrochemical reactor designed to produce acidic solutions as low as pH 1 for iCGP dissolution and alkaline solutions as high as pH 13 for acid neutralization and iCGP re-precipitation.

The electrochemical reactor consisted of an anode, middle, and cathode chamber – each holding 12 mL – secured by three hollow Perspex plates (5.3 × 10.2 × 1.2 cm³) and bolted between two solid two Perspex plates (10.1 × 1.3 × 8.7 cm3). A bipolar membrane (Fumasep FBM, Fumatech GmbH, Germany) separated the anode and middle chambers, while a cation exchange membrane (CMI-7000, Membranes International Inc., USA) separated the middle and cathode chambers. The anode was a titanium mesh coated with IrO₂/Ta₂O₅ (6 cm², Magneto Special Anodes, Netherlands), and the cathode was stainless steel (6 cm2, 316 stainless steel, Small Parts, Plymouth, MI). To assess acid and base production, pH was monitored in each chamber (FP220, Mettler Toledo, USA) under varying electrolyte concentrations (0.05, 0.1, and 1 M potassium salts). Potassium sulfate served as the anolyte due to its oxidative stability, while potassium chloride was used in the middle and cathode chambers for its resistance to reduction.

Potassium concentrations were verified via ion chromatography on a Dionex ICS-6000 (IC, Thermo Fisher/Dionex chromatograph, IonPac SCS1 column, unsuppressed, 4 mM tartaric acid and 2 mM oxalic acid eluent, 1.0 mL/min, 30 °C).

### 3.7 Electrochemical Cyanophycin Solubility Manipulation

In each test, 200 mL of electrolyte was circulated through the respective chambers (K2SO4 for the anode and KCl for the middle and cathode) under a constant current density of 10 mA/cm². Because electrolyte concentration influences solution conductivity and energy consumption (Apraku et al., 2024), we investigated the impact of electrolyte concentration on operational voltage and current via potentiostat (VMP-300, BioLogic Sciences Instruments, Seyssinet-Pariset, France) to compare energy consumption across concentrations.

For iCGP recovery, chronoamperometry (constant voltage of 5 V; 1 M potassium electrolyte) was used to generate highly acidic (pH 1) and basic (pH 13) solutions, because higher electrolyte concentrations facilitate greater pH shifts under these conditions. To dissolve iCGP, 100 mg of dried iCGP derived from 6308CphA1 was mixed into 20 mL of the acid solution and sonicated for 20 minutes. After complete dissolution, the solution was titrated to neutral pH (∼7) using the electrochemically generated base to precipitate iCGP. Precipitated iCGP was collected via centrifugation at 4,500 rpm for 25 min (Sorvall ST 8, Thermo Fisher, Waltham, MA) and dried in a vacuum oven at 60 °C overnight prior to gravimetric quantification.

Equation 1 defines the recovery efficiency *ε* as the ratio of the precipitated dry iCGP mass *M_f_* (g of iCGP) over the initial dissolved iCGP mass *M*_0_ (g of iCGP).

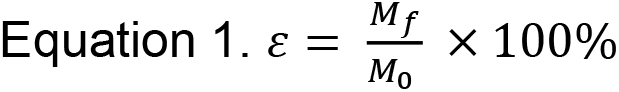

The specific energy consumption (MJ/kg iCGP) was calculated using Equation 2 to estimate the energy required per kilogram of iCGP recovered. This value reflects the energy input during acid and base production, calculated as the product of voltage (*V*), electrode area (*A*),conductivity (*σ*), and time (*t*), and is normalized by the initial dissolved iCGP mass (*M0*) and the recovery efficiency (*ε*), as defined by Equation 1. The calculation also accounts for the volume ratio of acid and base consumed relative to the stock solutions generated during chronoamperometry.

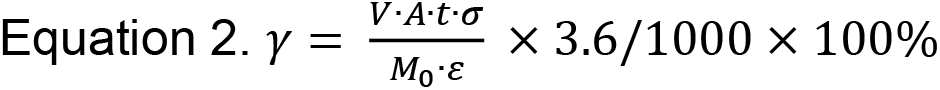

### 3.8 Cyanophycin Visualization via SDS-PAGE

Between approximately 5-10 mg cell wet weight was transferred to a pre-weighed 1.5 mL microcentrifuge tube and resuspended with premixed Takara Xtractor buffer (San Jose, CA) to a final concentration of 50 mg cell wet weight/mL. Cell lysis was performed via gentle inversion for 10 min at room temperature. Where specified, soluble and insoluble fractions of the lysate were separated via centrifugation at 17,000 x g. Supernatant was drawn off as the soluble fraction and the remaining solids were resuspended in an equal volume of water as the insoluble fraction. In preparation for gel loading, 20 μL of each sample was mixed with 20 μL 2x Laemli buffer (containing β-mercaptoethanol) in 200 μL PCR tubes and heated to 95 °C via thermocycler for 10-20 min. Sodium dodecyl sulfate-polyacrylamide gel electrophoresis (SDS-PAGE) was performed using precast 11.5% (w/v) polyacrylamide gels (Bio-Rad, Des Plaines, IL). Gels were run at 120 V for 60 min and subsequently treated with InstantBlue Coomasie Protein Stain (AbCam, Cambridge, UK) for at least 60 min prior to imaging.

### 3.9 Cyanophycin Titer Quantification via Sakaguchi Reagent

A fraction of the previously washed and resuspended cell pellet was transferred to a preweighed 200 μL PCR tube and incubated at 70 °C overnight. The difference in mass was used to calculate g CDW/mL washed cells. Arginine content was quantified as a proxy for cyanophycin as it has a higher arginine content (25-50 mol%) (Tseng et al., 2012) compared to bacterial proteins (3-7 mol%) (Brüne et al., 2018). Arginine in the acid extract supernatant was quantified using the Sakaguchi reagent as described previously but scaled to ∼200 μL per reaction for 96-well plate compatibility (Messineo, 1966). Samples were analyzed for endpoint absorbance at 520 nm for 10 min after reaction assembly. Arginine standards (0-100 μg/mL in 0.1 M HCl) were prepared and analyzed alongside samples on the same plate.

To account for background arginine contributions unrelated to cyanophycin — including heterologous CphA1 expression — a negative control strain expressing the red fluorescent protein mCherry but incapable of producing cyanophycin was cultivated and analyzed in parallel. The arginine content of this control strain was treated as baseline. For CphA1-expressing strains, the difference in arginine content relative to this baseline was attributed to cyanophycin. Assuming a defined mol% arginine in cyanophycin, this difference was used to estimate the per-cell cyanophycin titer.

Cyanophycin is canonically composed of 50 mol% arginine (Berg et al., 2000), corresponding to ∼56.7% (w/w). However, prior studies of heterologous CphA1 expression in *E. coli* have consistently reported lysine incorporation into the polymer (Tseng et al., 2012), (Frommeyer & Steinbüchel, 2013a). As lysine is thought to substitute for arginine rather than aspartic acid during polymerization, lysine-rich cyanophycin is commensurately arginine-poor. A cyanophycin sample with a reported lysine fraction of 25 mol% lysine, for instance, has an implied arginine content of 25 mol%. Accordingly, the estimated cyanophycin titer derived from such an assumed composition would be nearly twice that calculated using the 50 mol% assumption. In light of this variability, we therefore assumed the canonical 50 mol% arginine composition such that the cyanophycin titers derived from such measurements would conservatively represent the lower boundary of feasible values.

### 3.10 Optical Density and Fluorescence Measurements

Optical measurements were conducted via 96-well plates in a BioTek microplate reader. 300 μL per sample was measured for optical density via absorbance at 600 nm or mCherry fluorescence via excitation/emission at 585/615 nm, maximum gain.

### 3.11 Ammonium Quantification

Ammonium quantification was completed using HACH Ammonia TNTplus vial test kits and a HACH DR3900 spectrophotometer (Loveland, CO). Culture samples were clarified via centrifugation, diluted with water to the working range of 2-47 mg/L NH4-N and added to the reaction vial. Samples were incubated at room temperature for 15 min prior to reading.

## 4. Results

### 4.1 Estimating Cyanophycin Content by a Rapid Arginine quantification Method

Cyanophycin quantification is inherently challenging due to its amino acid composition, since arginine and aspartate are abundant intracellularly both as free metabolites and as components of cellular proteins. Consequently, amino acid analysis of whole-cell lysates cannot resolve cyanophycin content without prior separation. Established methods typically address this by exploiting cyanophycin’s pH-dependent solubility, which permits selective dissolution, re-precipitation, and subsequent gravimetric measurement (Frey et al., 2002). In practice, however, gravimetric measurement imposes constraints on culture scale, since adequate biomass is needed to obtain measurable amounts of polymer; recovery of 10 mg cyanophycin from *E.* coli containing 5% (w/w) cyanophycin, for instance, would require approximately 300 mL of culture at OD₆₀₀ ≈ 2 (0.33 g CDW/L), even without consideration for potential purification losses.

To overcome this limitation, we investigated the use of the Sakaguchi reagent to directly quantify arginine from the liquid-phase acid extract of cell pellets, thereby circumventing the need for cyanophycin precipitation and gravimetric analysis (Materials and Methods 3.9). While this reagent is well established for staining cyanophycin-containing cells (Watzer & Forchhammer, 2018), (Ke & Haselkorn, 2013), (Elbahloul et al., 2005), (Sukenik et al., 2015) or quantifying arginine in purified cyanophycin (Simon, 1971), its direct application to acid extracts, to the best of our knowledge, has not been reported. Furthermore, even when employed on purified samples, studies often omit reporting the arginine content of a cyanophycin-negative control, potentially confounding the measured signal with contributions from non-cyanophycin sources such as free arginine or native host proteins (Elbahloul et al., 2005), (Burgstaller et al., 2022).

To evaluate this potential bias, we cultivated a cyanophycin-negative *E. coli* strain expressing mCherry alongside a cyanophycin-producing strain expressing *Synechocystis* sp. 6308 CphA1 (6308CphA1) in LB supplemented with 10 mM L-arginine. After cells were washed, the acid extracts from the biomass of both strains were analyzed following qualitative confirmation that the 6308CphA1 strain produced a distinct polymer band consistent with cyanophycin (Fig. 1A). The appearance of protein products in the acid extract for the no-cyanophycin control (mCherry) demonstrates that the extraction process was not perfectly selective. Sakaguchi analysis revealed a substantial background arginine signal in this negative control, representing ∼44% of the signal detected in the 6308CphA1 extract (Fig. 1B). Cyanophycin content was estimated by subtracting the arginine in the no-cyanophycin control from the cyanophycin control and assuming an arginine composition of 50 mol% (56.7% w/w). Using this approach, the per-cell cyanophycin content of *E. coli* expressing 6308CphA1 was calculated as 1.57 ± 0.41% (g/g CDW).

### 4.2 Cyanophycin Synthetase Selection

We next applied our cyanophycin quantification method to evaluate three previously characterized cyanophycin synthetase homologs. Among them, *Synechocystis sp.* PCC 6308 *cphA1* (6308CphA1) is regarded as a benchmark enzyme and has been featured in numerous studies (Frey et al., 2002),(Voss & Steinbüchel, 2006). However, it is known to suffer from expression and solubility issues (Swain et al., 2023). Additionally, it has been shown to produce predominantly soluble cyanophycin (sCGP), reducing the potential efficacy of the proposed pH-based separation method which can only recover insoluble cyanophycin (iCGP).

To circumvent this, we tested two alternative synthetases, *Acinetobacter baylyi* ADP1 *cphA1* (AbCphA1) and *Tatumella morbirosei cphA1* (TmCphA1), which have been reported to produce predominantly iCGP (Swain et al., 2023),(Elbahloul et al., 2005). All three enzymes were active in LB medium supplemented with 10 mM L-arginine. AbCphA1 demonstrated the best performance in these conditions and achieved cyanophycin accumulation of 3.1 ± 1.5% CDW after 48 hours (Fig. 2). Interestingly, the two synthetases that favor insoluble product formation did not show accumulation until after the 21 hr time point. This delayed onset is consistent with the absence of a cyanophycinase-like domain that is present in 6308CphA1, but not AbCphA1 and TmCphA1 (Sharon et al., 2022). This cyanophycinase domain is hypothesized to create new primers upon which cyanophycin chains are extended by partially degrading existing cyanophycin, decreasing the lag time preceding robust cyanophycin accumulation. Despite the potential advantage, it is possible that this domain is also responsible for unintended cyanophycin degradation should the rate of decomposition ever exceed the rate of polymerization, which could occur during prolonged incubations (SI Fig. 3). For this reason, homologs with and without the domain, 6308CphA1 and AbCphA1, were selected for further evaluation using mock manure hydrolysate media.

### 4.3 Cyanophycin Engineered Strains Recover Nitrogen from Mock Manure Hydrolysates

Cyanophycin production was next evaluated in mock manure hydrolysate medium, formulated with glucose, xylose, arabinose, and amino acids as the primary carbon and nitrogen sources (Materials and Methods, 3.3). To probe the impact of nitrogen source on cyanophycin recovery, a secondary, high ammonium condition was simulated by replacing approximately 60 mM casamino acids (6.92 g/L) with 60 mM ammonium sulfate (7.93 g/L). Cultures expressing either 6308CphA1 or AbCphA1 began accumulating cyanophycin within 48 hrs (SI Fig. 4). After 144 hrs, cultures were harvested and cyanophycin content was quantified. Compared to 6308CphA1, AbCphA1 exhibited enhanced cyanophycin accumulation under high ammonium conditions, yielding the highest per-cell titer (2.1 ± 0.3%, Fig. 3B) and volumetric titer (70.2 ± 5.3 mg/L, Fig. 3C) despite a comparatively lower optical density. Whereas biomass-normalized cyanophycin titers for 6308CphA1 remained consistent across nitrogen sources, AbCphA1 exhibited a five-fold enhancement under organic nitrogen compared with high ammonium conditions. Under high ammonium conditions, the 6308CphA1-expressing strain exhibited significantly greater specific ammonium uptake compared to the mCherry control (Fig. 3D), consistent with the putative role of cyanophycin as a nitrogen-sink.

**Figure 4.**
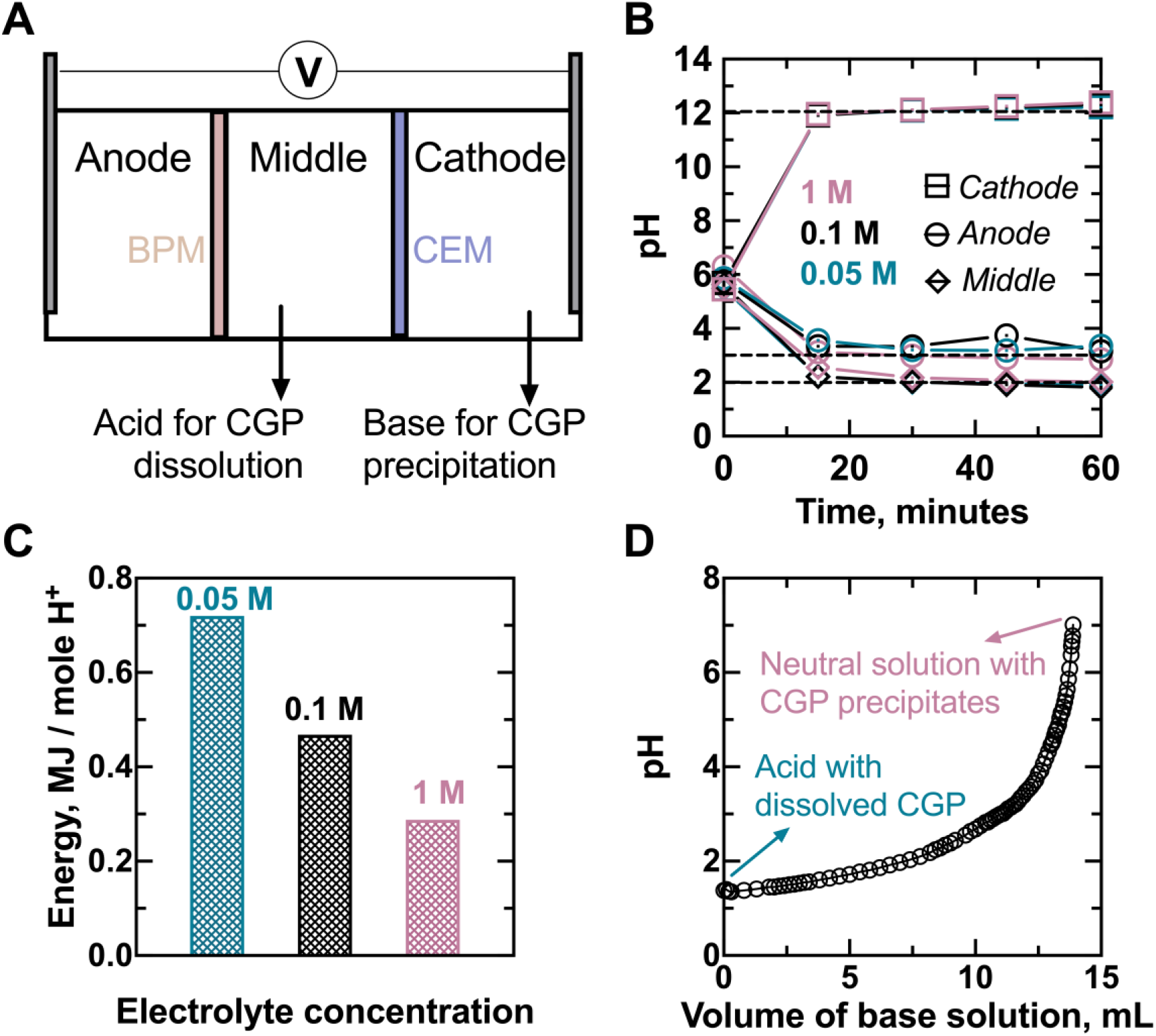
Electrochemical acid and base production enables cyanophycin separation. (A) Electrochemical reaction schematic for acid and base production. (B) pH changes in different chambers of the electrochemical reactor with different electrolyte concentrations (0.05, 0.1, and 1 M as potassium; K2SO4 was used as anolyte, and KCl was used as the electrolyte in both the middle and cathode chamber). (C) Energy consumption per mole of protons produced at different electrolyte concentrations. (D) Titration curve for 100 mg iCGP dissolved in acid solution (middle chamber electrolyte, 1 M KCl at pH 1.4) titrated with electrochemically produced base (1 M KCl catholyte at pH 13). Acid and base were both produced at 5 V. BPM, bipolar membrane; CEM, cation exchange membrane

### 4.4 Electrochemical Acid and Base Production Allows Sustainable Cyanophycin Separation

The separation of cyanophycin from background biomass requires strong acids and bases, incurring costs related to the production, transportation, and storage of hazardous chemicals. We overcame these challenges by designing an electrochemical reactor (Fig. 4A) to enable on-demand acid and base production using electricity and evaluated the reactor’s acid/base generation, energy consumption, and cyanophycin recovery efficiency.

Acid and base production was first assessed by monitoring pH changes in each reactor chamber using a 0.05 M electrolyte solution (0.05 M K₂SO₄ as anolyte; 0.05 M KCl in the middle and cathode chambers) under a constant current density of 10 mA/cm². Within one hour, mild acid was produced in both the anode (from initial pH 6 to final pH 3) and middle chambers (from pH 6 to 2), while base was produced in the cathode chamber (from pH 6 to 12), demonstrating successful electrochemical generation. The pH drop in the anode was attributed to the H+ generation during the oxygen evolution reaction, while the lower pH in the middle chamber resulted from water dissociation within the bipolar membrane. Because water dissociation proceeds without gas evolution, it is considered a more efficient acid-generating mechanism than oxygen evolution, consistent with the observed lower pH in the middle chamber. The pH increase in the cathode was caused by the OH-via the hydrogen evolution reaction.

Given that electricity is the primary input for this process, we next evaluated energy consumption across different electrolyte concentrations. We varied electrolyte concentration instead of voltage or current, which previous studies have shown only modestly affect pH outcomes on timescales relevant to cyanophycin extraction (Apraku et al., 2024), (Dong et al., 2022). Although pH changes were similar across concentrations (Fig. 4B), energy consumption was nearly halved at 1 M compared to 0.05 M (Fig. 4C). Based on these findings, we selected 1 M electrolyte for further experiments to generate stronger acid (pH ∼1) and base (pH ∼13) solutions at 5 V (current density ∼25 mA/cm²) for iCGP separation.

Finally, to test the efficacy of the produced chemical regenerant in modulating cyanophycin solubility, iCGP derived from 6308CphA1 was added *ex-situ* to acid from the middle chamber. Following dissolution, base from the cathode was used to titrate the solution to pH 7 and precipitated iCGP was measured gravimetrically. The titration curve showed a nonlinear pH increase, suggesting a buffering effect by the dissolved iCGP (Fig. 4D). Nevertheless, the required volume of base (∼15 mL) was less than that of acid (∼20 mL), enabling separation using a single batch of acid/base production with equal electrolyte volumes in each chamber. The overall iCGP recovery efficiency was 53.3% ± 4.2%, which, while improvable, demonstrates a proof of concept for electrochemical CGP separation via in situ acid/base generation.

### 4.5 Cyanophycin Synthetase N-Terminal Domain Influences CphA1 Expression and Cyanophycin Solubility

To enhance overall cyanophycin production and increase the fraction recoverable via pH manipulation (iCGP), we evaluated the impact of the CphA1 N-terminal domain on enzyme expression and cyanophycin solubility. In both LB and mock manure hydrolysates, SDS-PAGE of whole-cell lysates revealed peptide aggregates of approximately 100 kDa, consistent with the size of individual CphA1 subunits (Fig. 2, SI Fig. 4). The relative intensity of CphA1 bands, compared to the highly expressed and soluble control protein mCherry, indicated disproportionate accumulation of CphA1. Fractionation of whole-cell lysates further demonstrated that these aggregates were predominantly insoluble (Fig. 5A). The formation of insoluble, inactive inclusion bodies is a well-documented outcome of heterologous protein expression (Bhatwa et al., 2021), and difficulties in expressing 6308CphA1 have been reported previously (Swain et al., 2023). Consistent with this, early experiments varying the expression vector, promoter, and ribosome binding site produced substantial changes in both apparent CphA1 aggregation and cyanophycin production (SI Fig. 5). Because the N-terminus emerges first from the ribosome, it has been shown to strongly influences peptide folding and solubility in a broad range of proteins (Kim et al., 2007), motivating our subsequent focus on this domain.

**Figure 5.**
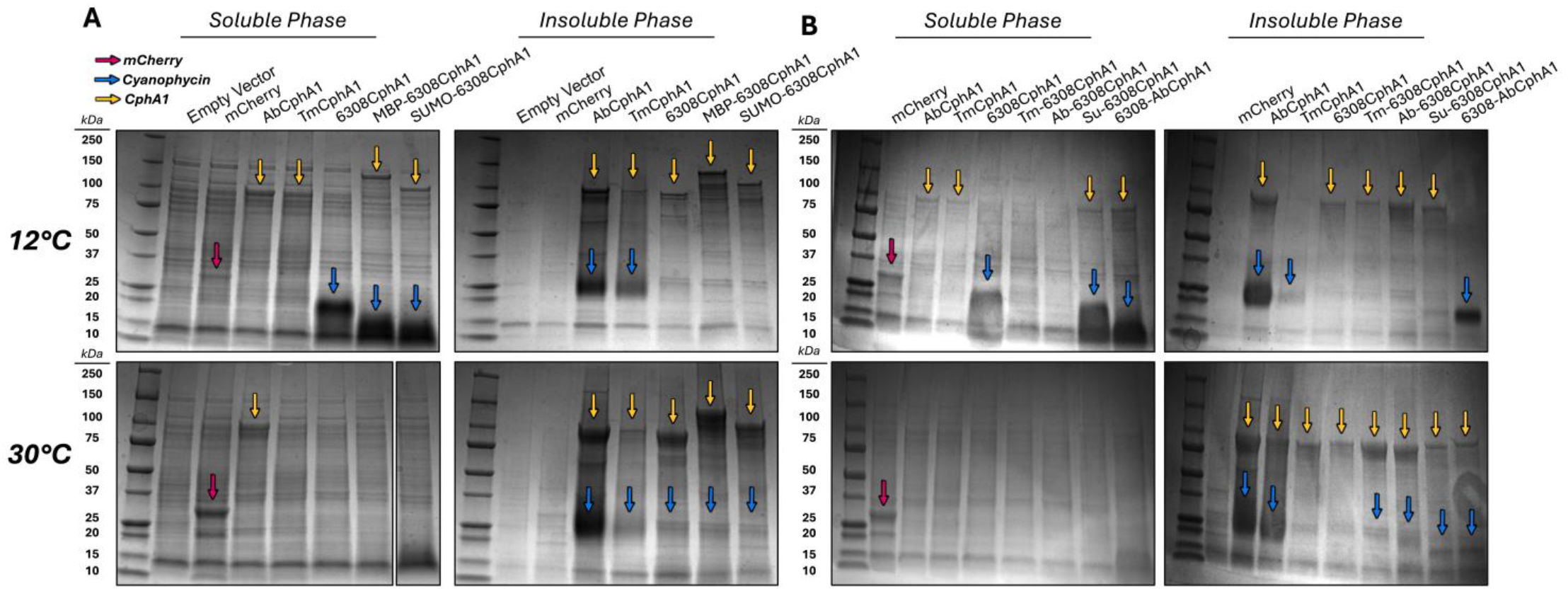
Impact of N-terminal modifications on CphA1 expression and cyanophycin solubility at different cultivation temperatures. (A) SDS-PAGE gel of soluble versus insoluble fractions of cells expressing different CphA1 homologs including those with solubility tags. One lane was excised from the soluble phase, 30 °C gel for illustrative simplicity. (B) SDS-PAGE gel of soluble versus insoluble fractions of cells expressing CphA1 chimeras. Cultures were grown in unsupplemented LB medium for 24 hrs prior to harvest.

To address the challenge of insoluble CphA1 accumulation, we tested two strategies: fusion of solubility-enhancing tags and substitution of the N-terminal domain between CphA1 homologs. We first fused two commonly used solubility tags, maltose binding protein (MBP) and small ubiquitin-like modifier (SUMO), to the N-terminus of 6308CphA1. Although both fusions produced cyanophycin at 12 °C, neither produced obvious increases in CphA1 solubility or cyanophycin production at 30 °C, a temperature more relevant to process-scale cultivation (Fig. 5A). Notably, both fusions produced soluble cyanophycin with a lower molecular weight than the wild-type protein, suggesting a possible enhancement of cyanophycinase-like activity from the N-terminal domain.

Given the lack of improvement in CphA1 solubility at 30 °C we next considered whether poor expression might be linked to improper folding of the N-terminal domain itself during high-level expression. The apparent solubility of unmodified CphA1 homologs varied by temperature: AbCphA1 was largely detected in the insoluble fraction at both 12 °C and 30 °C, 6308CphA1 appeared soluble at 12 °C but not at 30 °C, and TmCphA1 showed only a small insoluble fraction under both conditions but coincided with lower levels of cyanophycin (Fig. 5A/B). To explore whether these solubility patterns were influenced by the N-terminal domain, we generated chimeric variants by swapping domains between homologs. In support of this hypothesis, the 6308-AbCphA1 chimera showed a solubility profile more similar to 6308CphA1, with a smaller insoluble fraction at 12 °C, while Ab-6308CphA1 resembled AbCphA1, with a larger insoluble fraction at the same temperature (Fig. 5B). This pattern, however, was not consistent across all constructs: Tm-6308CphA1 displayed increased aggregation at 30 °C unlike the unmodified TmCphA1. While it is possible that this aggregation was caused by incompatibilities between the chimeric domains, the production of cyanophycin by Tm-6308CphA1 demonstrates that any such effect was not catastrophic. More likely, the N-terminal domain is not always a key determinant of CphA1 expression. Interestingly, while 6308CphA1 appeared to produce predominantly sCGP and AbCphA1 predominantly iCGP, the 6308-AbCphA1 chimera produced both forms. These results implicate the N-terminal domain in influencing cyanophycin solubility.

## 5. Discussion

This study constitutes a proof of concept for an integrated nitrogen-recovery bioprocess at manure generation sites. We established a workflow for cyanophycin biosynthesis and recovery from mock manure derivatives and applied electrochemical methods to enable simple, low-input purification of cyanophycin. Our results reveal key challenges and highlight opportunities for improving both biosynthesis and downstream processing of this nitrogen-rich biopolymer.

### 5.1 Cyanophycin Synthetase Expression Remains a Critical Parameter to be Optimized

Cyanophycin production was compared among three cyanophycin synthetase homologs: 6308CphA1, AbCphA1, and TmCphA1. Based on arginine measurements and qualitative SDS-PAGE analysis, cyanophycin accumulation broadly followed the trend 6308CphA1 ≈ AbCphA1 > TmCphA1, contrasting with previous reports in which TmCphA1 exhibited more than twice the specific activity *in vitro* (Sharon et al., 2021), (Krehenbrink & Steinbüchel, 2004), (Aboulmagd et al., 2001) and produced more than twice the cyanophycin *in vivo* (Swain et al., 2023) compared to 6308CphA1 and AbCphA1. While this study employed LB rather than TB medium as used previously for these comparisons, the consistent underperformance of TmCphA1 across experiments indicates that intrinsic enzyme activity alone is an insufficient predictor for cyanophycin production, underscoring the importance of cellular context and expression conditions in shaping *in vivo* performance.

To deconvolute the many factors likely influencing the functional activity of expressed CphA1 homologs, we examined the presence of insoluble inclusion bodies by SDS-PAGE. Although inclusion body formation is often associated with loss of enzyme function (Bhatwa et al., 2021), their presence did not always preclude cyanophycin production, nor did their absence always correlate with accumulation (Fig. 5, SI Fig. 4 and 5). Interestingly, while previous studies reported an increase in the specific activity of purified 6308CphA1 at elevated temperatures *in vitro* (Aboulmagd et al., 2001), cyanophycin production in this study was more consistent at 12 °C than at 30 °C. Although supported by the reduction in insoluble CphA1 aggregation at 12°C, this pattern may not reflect the impact of temperature on CphA1 solubility alone. Reduced growth rates at lower temperatures may have contributed to higher apparent per-cell cyanophycin titers by reducing the effect of dilution due to cell division.

Together, these findings suggest that *in vivo* cyanophycin accumulation is more significantly constrained by the complex interplay between enzyme expression, growth conditions, and host physiology, rather than by the intrinsic catalytic turnover rate of a given CphA1 homolog. Further investigation and potential engineering of a soluble variant with tailored compatibility for heterologous expression may therefore provide a promising avenue to enhance cyanophycin yields.

### 5.2 Recovery Efficiency is Contextually Dependent on Cyanophycin Concentration and Solubility

Considering the cost of strong acids and bases, the large volumes of saline waste and greenhouse gas emissions associated with their industrial production, and the logistical constraints of transporting them to potentially decentralized nitrogen waste generation sites, electrochemical acid and base generation was evaluated for its ability to support a pH-based purification strategy for 6308CphA1-derived iCGP (Kavvada et al., 2017). Building on previous work that evaluated the effects of current and voltage on electrical resistance and adsorptive ammonia recovery (Dong et al., 2022), our study focused on adjusting electrolyte concentrations to reduce energy consumption while maintaining effective acid/base strength. While successful in generating strong acids and bases, the moderate yields of recovered iCGP emphasize the need for improvement at the junction of cyanophycin production and recovery strategies, particularly since many factors appear to influence precipitation behavior.

The amino acid composition of cyanophycin, and consequently its solubility, varies across CphA1 homologs (Frommeyer & Steinbüchel, 2013a). Since the iCGP used was synthesized from 6308CphA1, it is likely that the lysine content was non-zero (Tseng et al., 2012), potentially reducing the polymer’s intrinsic tendency to precipitate. It is important to recognize that the classification of cyanophycin as “soluble” or “insoluble” is not absolute but rather contextually dependent on conditions such as pH and concentration. Like many biopolymers, cyanophycin exhibits a defined solubility limit at a given pH (e.g., pH 7), above which excess material will precipitate out. If the total cyanophycin concentration in the electrolyzer is too low, a significant fraction may remain in solution at this saturation threshold, resulting in lower apparent recovery. This limitation may be mitigated by reducing the extraction volume relative to the mass of cyanophycin, thereby increasing its concentration during neutralization and promoting precipitation. However, this strategy must be carefully balanced to avoid exceeding cyanophycin’s solubility limit under highly acidic or alkaline conditions, which would reduce overall extraction efficiency. Importantly, beyond losses due to solubility limits, this approach relies on the ability of cyanophycin to precipitate at a specific pH, rendering it unsuitable for recovering sCGP.

While sCGP is characterized by elevated lysine content (Tseng et al., 2012),(Frommeyer & Steinbüchel, 2013b), lysine itself is a high-value, essential amino acid for swine and poultry feed and significantly enhances the potential utility of cyanophycin as a feed additive (Liao et al., 2015). An optimal process would thus seek to maximize the amount of insoluble, lysine-rich cyanophycin generated. While these characteristics are not generally thought of as compatible, our work strengthens the claim that cyanophycin solubility is influenced by both amino acid composition as well as N-domain activity, which we hypothesize mediates solubility by modulating polymer size.

Cyanophycin produced by 6308CphA1, which is predominantly soluble, tends to have lower molecular weight than that produced by AbCphA1 (Fig. 2, Fig. 5). More broadly, all insoluble cyanophycin samples we examined were the same size or larger than their soluble counterparts, with one notable exception: the insoluble fraction of the 6308-AbCphA1 chimera. This iCGP was the only sample we observed with a molecular weight below 20 kDa. This result supports the hypothesis that a catalytically active N-terminal domain reduces cyanophycin chain length, thereby impairing the formation of insoluble granules. Previous work places the N-terminal domain as loosely binding the elongating cyanophycin strand rather than the amino acids to be incorporated (Sharon et al., 2021), suggesting that the lysine content of 6308CphA1-derived cyanophycin is likely independent from N-domain activity. Future genetic engineering efforts for this application should focus on inactivating the cyanophycinase-like activity of the 6308CphA1 N-terminal domain to increase polymer size and insolubility while preserving its favorable lysine incorporation profile. Notably, cyanophycin synthesized by a different *Synechocystis sp.* CphA1 (PCC 6803) has been reported to reach up to 100 kDa in size in cyanobacteria and plants, a differential attribute that remains unexplained in heterotrophs (Ziegler et al., 1998).

### 5.3 The Nitrogen Removal and Recovery Efficiency of the Proposed Process Rivals Those of Alternatives

Our overall goal is to demonstrate the feasibility of microbial nitrogen concentration and electrochemical recovery of cyanophycin. The efficacy of nitrogen recovery from the substrate was assessed by comparing the amount of nitrogen sequestered per gram of cell dry weight in our engineered strains. In the high ammonium formulation, the strain that could not synthesize cyanophycin (mCherry control) sequestered 90.6 ± 4.9 mg NH₄⁺-N/g CDW, whereas the 6308CphA1 and AbCphA1 strains sequestered 128.5 ± 9.4 mg NH₄⁺-N/g CDW and 113.8 ± 19.0 mg NH₄⁺-N/g CDW, representing 42% and 26% improvements over baseline, respectively. On an absolute basis, the 6308CphA1 strain removed the most nitrogen (512 mg NH₄⁺-N/L), although this accounted for only 27% of the initial NH₄⁺-N in the medium. While the timeframe of this experiment was ultimately constrained by volume losses from sampling and evaporation, the growth trajectory illustrated in SI Fig. 6 indicates that further growth and subsequent nitrogen sequestration would likely have continued after 144 hrs.

Future work should focus on optimizing the carbon to nitrogen ratio to maximize nitrogen removal per gram of manure-derived carbon. In addition, optimization of cultivation temperature to balance per-cell cyanophycin yield with total biomass yield could increase both overall nitrogen removal and recovery metrics.

Our energy analysis of the electrochemical recovery process revealed a consumption of approximately 15 MJ/kg iCGP, or about 65 MJ/kg nitrogen, assuming a nitrogen content of 23% in cyanophycin (50 mol% Arg). This places our method within the range reported for comparable electrochemical nitrogen separation techniques, such as electrochemical stripping (30–300 MJ/kg N)(Liu et al., 2020),(Tarpeh et al., 2018) and bipolar membrane electrodialysis with membrane contactors (BMED-MC, 10–100 MJ/kg N)(Li et al., 2021). Although we have not estimated energy costs associated with bacterial accumulation of cyanophycin, we anticipate they would be modest. This result suggests that electrochemical cyanophycin recovery may be competitive with other nitrogen recycling approaches, especially if integrated into renewable energy-driven bioprocesses. Continued optimization of the purification process could further reduce energy requirements by identifying extraction modes that use less acid and base per sample, such as using higher biomass concentrations per extraction or raising the pH setpoint for acid extraction. In parallel, control variables such as extraction temperature and sample agitation could be evaluated for potential tradeoffs between efficacy and energy consumption.

### 5.4 Outlook

In this work, we demonstrate a proof of concept process for biological nitrogen removal and recovery at potentially decentralized sites of waste generation using genetically engineered *E. coli* and electrochemical acid/base generation to enable on-site production and purification of cyanophycin. Our results indicate that the final output, yield of recovered cyanophycin, depends heavily on both intracellular titer and the relative insolubility of the polymer. These factors are in turn influenced by the interactions between chassis organism physiology, CphA1 homolog activity and expression, cultivation temperature, and media composition. Enhancing the production and recovery of iCGP will be essential for improving overall process feasibility and enabling scalable implementation. Furthermore, it is necessary to demonstrate that an inoculum of engineered *E. coli* can successfully metabolize nitrogen in non-sterile manure containing an existing microbial community. Such trials may face challenges with respect to robust *E. coli* growth as a consequence of competing microorganisms, inhibitory metabolites, and manure heterogeneity not accounted for with the sterile, defined mock medium employed here. Given the breadth of recombinant organisms previously modified to produce cyanophycin, consideration of alternative engineered hosts or communities capable of accumulating cyanophycin may prove fruitful for robust deployment in medium derived from manure hydrolysates. That said, success in this arena would establish an effective recycling stream within the synthetic nitrogen cycle, decreasing reliance on single use nitrogen inputs and providing animal farmers with a more consistent approach to waste management in the face of an ever-changing regulatory landscape.

## Supporting information

DNA Sequences

## Acknowledgements

We would like to thank Prof. Marting Shmeing for providing the codon-optimized nucleotide sequence for TmCphA1 and for his assistance in choosing a split site for the chimeric mutants.

## Author Contributions: CRediT

**Kevin Fitzgerald**: conceptualization, data curation, investigation, methodology, visualization, writing – original draft, review, and editing; **Hang Dong**: investigation, methodology, data curation, visualization, writing – original draft; **Edward Apraku**: investigation, methodology, data curation, visualization, writing – editing; **Md Aminul Islam Prodhan**: conceptualization, investigation, resources, writing – original draft; **Dylan Hakken**: investigation; **George Wells**: conceptualization, project administration, resources, supervision, writing – editing; **William Tarpeh**: conceptualization, project administration, resources, supervision, writing – editing**; Keith Tyo**: conceptualization, project administration, resources, supervision, writing – editing.

## Declaration of AI Use in the Manuscript Preparation Process

During the preparation of this work, the authors used ChatGPT services to aid in the writing process to improve sentence structure and readability. After using this tool, the authors reviewed and edited the content as needed and take full responsibility for the content of the published article.

