## Supplementary Information for "Demonstration of an Integrated Process for Manure-Based Nitrogen Recovery Using the Biopolymer Cyanophycin"

Corresponding author: Keith Tyo

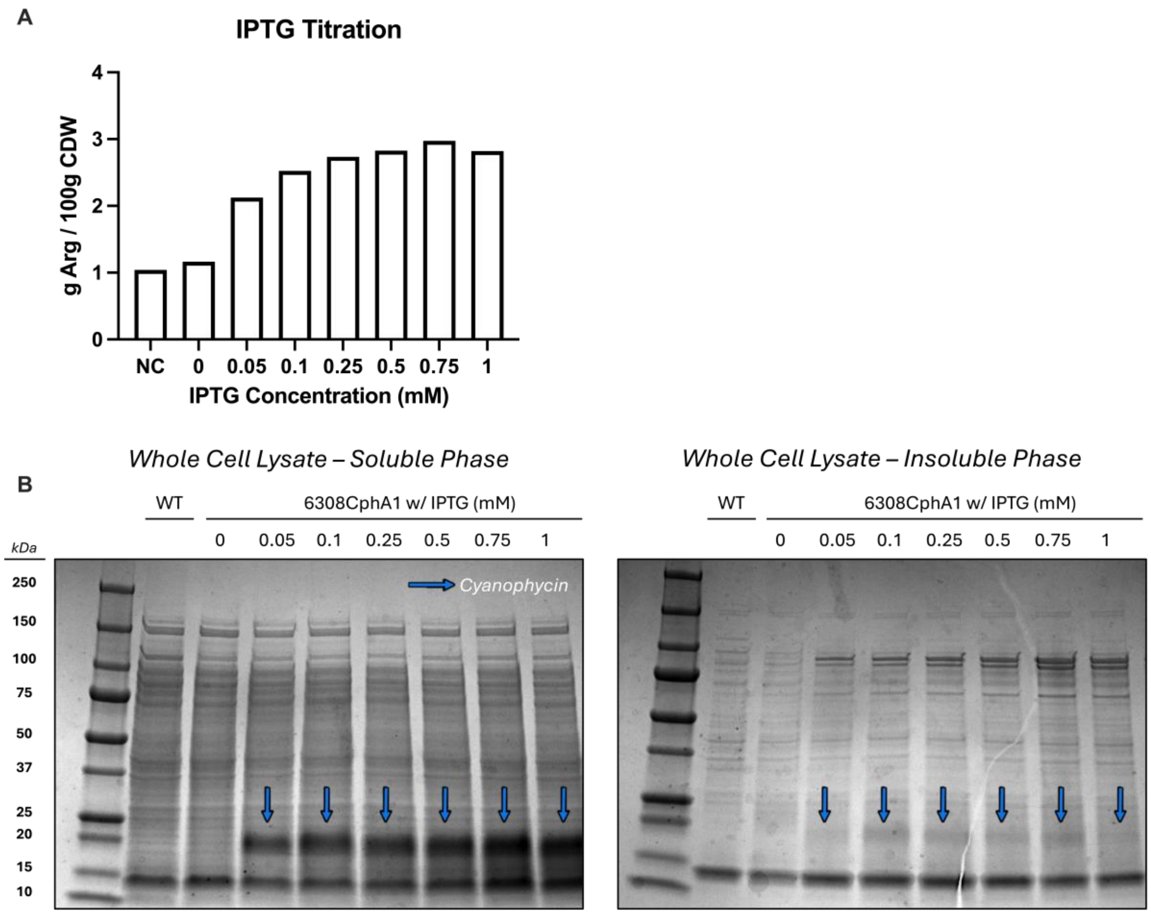


**Supplementary Figure 1. IPTG Titration Curve for 6308CphA1.** *E. coli* BL21(DE3) pLysE was cultured at 12 °C in TB medium and induced with various concentrations of IPTG at a target $OD_{600}$ = 0.5. 0.75 mM IPTG correlated with the highest cyanophycin content following Sakaguchi quantification in agreement SDS-PAGE results. Virtually all cyanophycin appears in the soluble fraction.

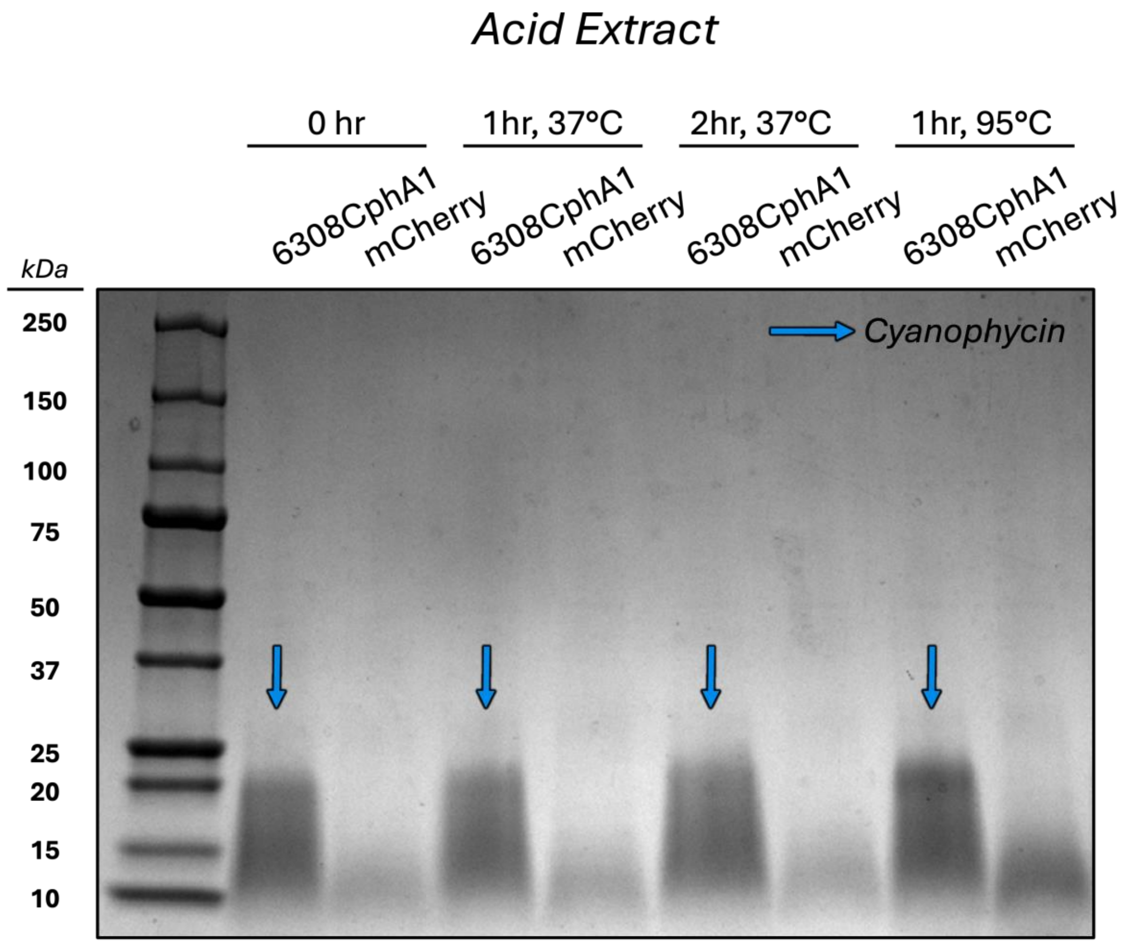


**Supplementary Figure 2. Optimization of Acid Extraction Procedure.** Different extraction regimes for 6308CphA1-derived cyanophycin were tested to determine the optimal conditions for maximizing cyanophycin extraction and minimizing non-specific peptide extraction. mCherry represents the cyanophycin-negative control. Notably, the amount of cyanophycin extracted did not seem to visibly increase with an increase of incubation time at 37 °C. While cyanophycin extraction efficiency appears to be enhanced at 95 °C, a concurrent increase in background peptide from the mCherry control was also observed.

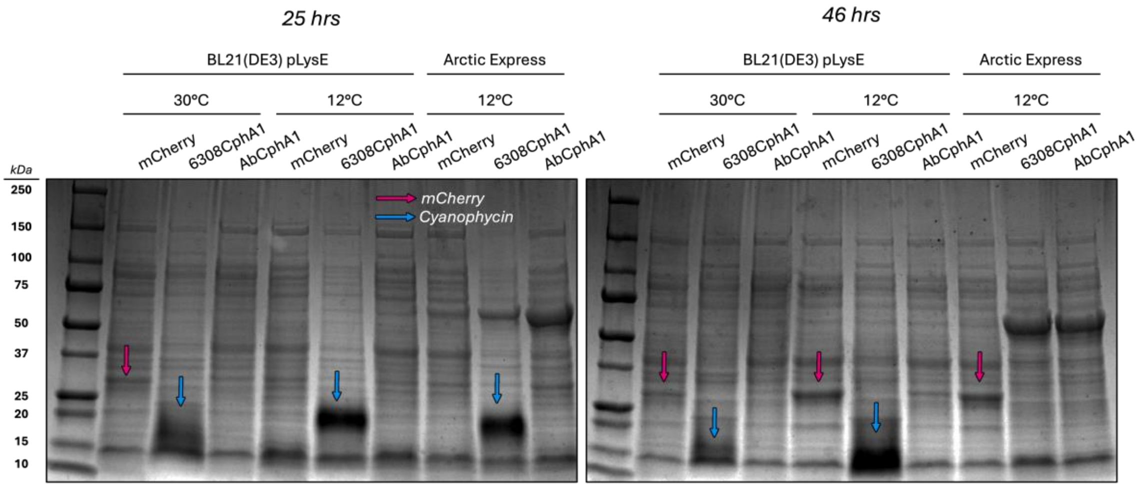


**Supplementary Figure 3. Comparison of *E. coli* Strains and Growth Temperatures.** To evaluate whether reduced cyanophycin yields at 30 °C were due to improper enzyme folding, *E. coli* BL21(DE3) pLysE was compared to *E. coli* ArcticExpress — a strain engineered for low-temperature expression — for cyanophycin production. Both strains were cultured at 12 °C, the recommended temperature for ArcticExpress. Cyanophycin production was detectable in both strains under these conditions, supporting the use of 12 °C for subsequent experiments. A decrease in cyanophycin molecular weight was observed from day one to day two in both strains, with polymer levels appearing to diminish entirely in ArcticExpress.

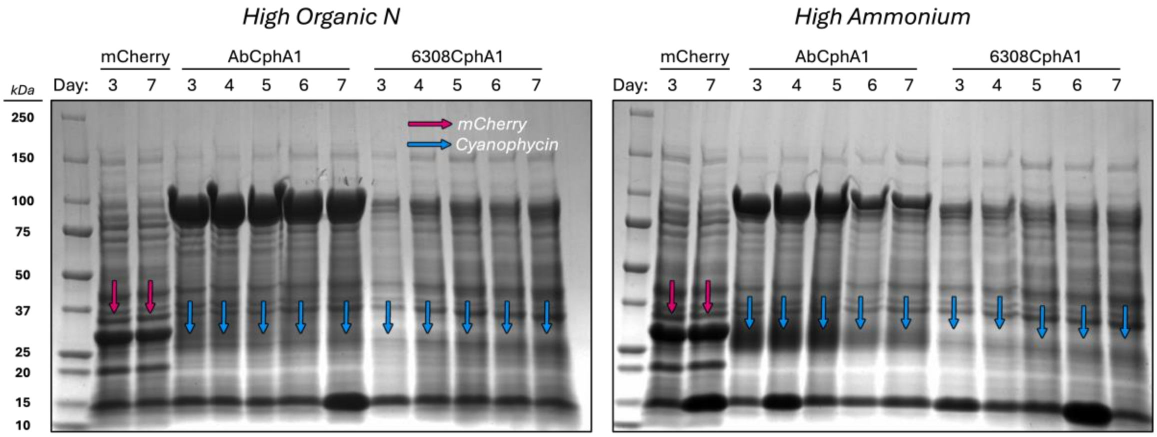


**Supplementary Figure 4. Time-Course Cyanophycin Production in Mock Manure Hydrolysate Media*.***

Samples from a single replicate were processed and visualized via SDS-PAGE.

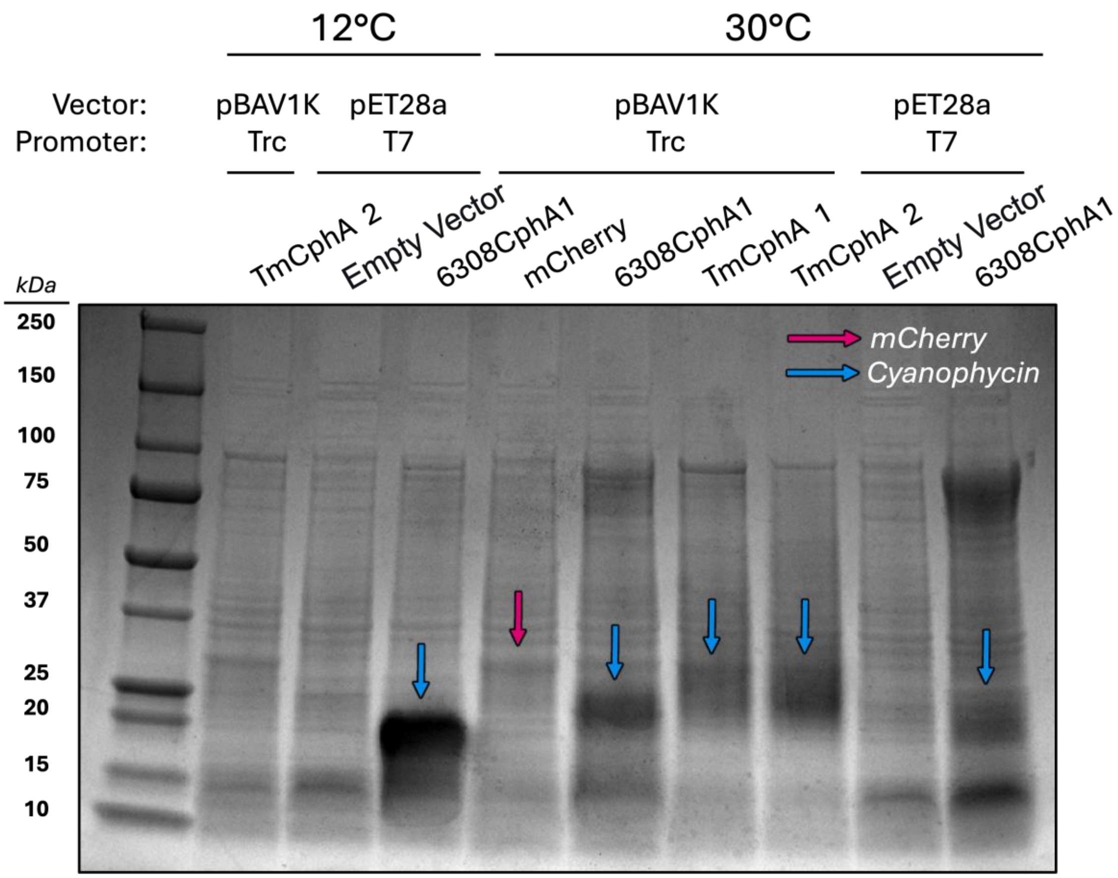


**Supplementary Figure 5. Impact of Expression Vector on CphA1 Expression and Cyanophycin Synthesis.** Both Trc and T7 promoters are induced with IPTG but vary in strength of expression. Expression of 6308CphA1 from pBAV1K-Trc appears to correlate with enhanced cyanophycin accumulation and reduced CphA1 aggregation compared to pET28a-T7 at 30 °C.

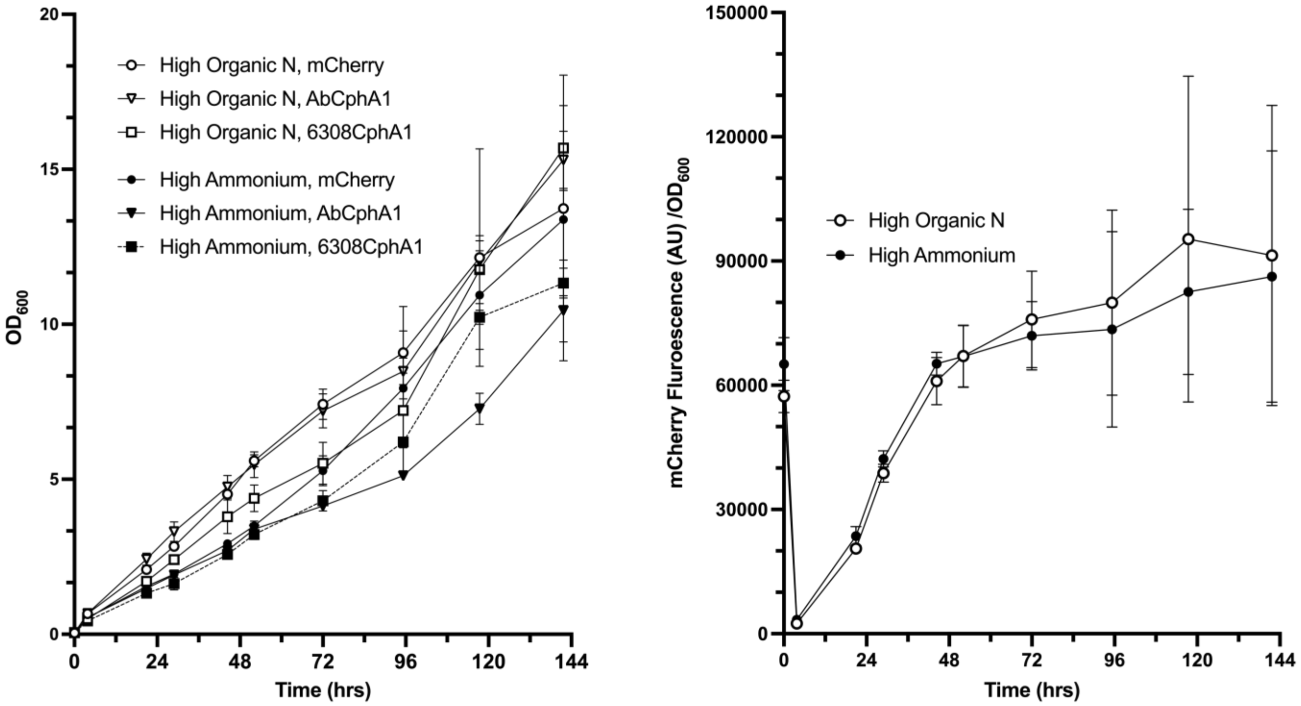


**Supplementary Figure 6. Optical Density and Specific mCherry Fluorescence on Mock Manure Hydrolysate Media.** Error bars = SEM for biological triplicate.
